# The use of high-resolution computed tomography to explore age-related trabecular change in human ribs

**DOI:** 10.1101/2021.07.12.452053

**Authors:** Sophia R. Mavroudas, Victoria M. Dominguez

**Affiliations:** Texas State University, San Marcos, TX 78666, USA; Dept. of Anthropology, Lehman College, City University of New York, Bronx, NY 10468, USA; Dept. of Anthropology, The Graduate Center, City University of New York, New York, NY 10016, USA; New York Consortium of Evolutionary Primatology, New York, NY, USA

**Keywords:** Rib, Age-at-death, Trabecular analysis, Bone histology, Forensic anthropology, microCT

## Abstract

High-resolution computed tomography was used to explore age-related trabecular change in male human ribs ranging in age from 20–95 years (Mean=55 years, SD=21.634 years) from the Texas State Donated Skeletal Collection (TXSTDSC). Two regions of interest (ROIs), midshaft (50%) and anterior (75%) were extracted from each scan to analyze age-related trabecular change. Dragonfly V4.1 was used to isolate cortical bone volumes of interest (VOIs) and three trabecular VOIs for each ROI; one each along the cutaneous cortex, the center of the medullary cavity, and the pleural cortex. Each trabecular VOI was analyzed for bone volume fraction (BV/TV), trabecular thickness (TbTh), trabecular spacing (TbSp), connectivity density (Conn.D), and degree of anisotropy (DA), within and between the 50 and 75% ROIs. Overall, the cutaneous VOIs at both the 50% and 75% ROIs exhibited greater BV/TV, TbTh, and Conn.D when compared to the center and pleural VOIs. All results are consistent with expected biomechanical strain on human ribs. Both trabecular variables and cortical bone volume are only weakly associated with age. These results show that 3D analysis of trabecular bone volume does not improve visualization or understanding of trabecular bone changes with age over traditional 2D methods. Future research should incorporate female samples to explore sex-related trabecular change variation.

**HIGHLIGHTS:**

- 3D analysis of change in trabecular structure along the rib length and with age
- Trabecular spacing at midshaft shows highest correlations with age-at-death
- Relative cortical area is more strongly correlated with age in anterior ribs than at midshaft

## Introduction

Existing histological rib methods for estimating age-at-death rely primarily on the analysis of human cortical bone cross-sections under light microscopy (1–5), although recent efforts raise the question of incorporating trabecular bone in these assessments to improve reliability and repeatability (6). Despite this suggestion, trabecular bone’s relationship to age in the rib, unlike cortical bone (7, 8), has not been extensively examined. The aim of this research is to quantify 3D trabecular bone changes in the human 6th rib as it relates to age using micro-computed tomography. As the field moves towards reimagining histological age-at-death estimation methods beyond linear regression models (5, 9), a better understanding of the relationship between trabecular bone and age can help to determine the utility of using trabecular bone variation, rather than or in addition to traditional cortical bone variables, for improving the accuracy and repeatability of histological age-at-death estimates in the rib.

In terms of bone tissue, the primary differences between cortical and trabecular bone are the result of structural organization rather than material composition (10). Cortical bone consists of densely packed layers of lamellae that form the thick outer shell of bones inside which trabecular bone resides. This trabecular bone is composed of a network of narrow struts and plates oriented in different directions often described as “spongy” in appearance (10). These differences in structural organization are related to skeletal remodeling, the process of removing bone for maintenance or repair and reforming new bone in its place, which results in the consistent formation of quantifiable structures within the cortex known as secondary osteons. As we age, the number of secondary osteons increases and the quantification of these structures relative to cortical bone area generates a density measure that can be used to reliably estimate age-at-death (1, 11, 12). In trabecular bone, remodeling occurs via the formation of hemiosteons rather than secondary osteons (13), although neither these hemiosteons nor the bone area of the trabecular struts is typically considered in histological age estimation methods.

Instead, cortical bone observation has been favored for age-at-death analysis; however, the distinction between cortical and trabecular bone is rarely as straightforward as suggested by these methods and is in many cases arbitrary (14). How researchers choose to delineate the transition zone between trabecular and cortical bone can have serious implications for the resultant data. For example, including all of the trabecularized region along the endosteal border often inflates cortical area measures, artificially decreasing osteon densities, while excluding trabecularized regions deliberately ignores the remodeling activity those portions of the cortex represent (15). This can be a major issue when working with osteoporotic individuals (16). To resolve this issue, some researchers propose sidestepping it entirely by including trabecular bone when measuring cross-sectional area, generating a total bone area rather than just a measure of the cortex (6). This merits further exploration due to its potential for improving histological age estimates, particularly in older individuals. Thus far, rib trabecular quantification in histological sections has been limited since preservation and quantification of trabecular bone using 2D histological techniques is delicate and time-consuming work and existing samples are small. Recent studies demonstrate that significant patterns of trabecular loss with increasing age exist (6, 17) and that there are sex-specific differences, with females exhibiting accelerated loss compared to males (18).

While providing important data related to trabecular bone loss with age, these studies would benefit from a more thorough understanding of the 3D organization of trabecular bone within the rib. Although histological analyses rely on stereological principles for the 2D analysis of 3D structures, trabecular bone properties are more completely visualized in 3D space, due to the 360° plate and strut structure. Through micro-computed tomography, the 3D measures of trabecular change such as trabecular thickness, strut connectivity, and overall directionality (anisotropy) might contribute more to our understanding of how trabecular change occurs over time. To date, most 3D approaches examining bone microstructure focus on the gain or loss of cortical bone (8, 19), likely due to the limited resolution of technologies such as clinical computed tomography (CT). MicroCT, on the other hand, demonstrates imaging resolution sufficient to quantify the number and thickness of trabecular struts within various skeletal elements (20). The purpose of this study is twofold: (1) to explore the use of 3D microCT to provide a more complete picture of trabecular bone change with age in the rib and (2) to examine the utility of incorporating 3D trabecular analysis for estimating age-at-death.

## Materials and Methods

This sample consists of left or right middle ribs from N=40 males from the modern Texas State Donated Skeletal Collection (TXSDTDSC) housed at the Forensic Anthropology Center at Texas State (FACTS). Ribs were collected from deceased individuals ranging in age from 20– 95 years (Mean=55 years, SD=21.634 years). Each donor was received at FACTS between 2008 and 2018. Only male ribs were used to maximize sample size while controlling for the potential effect of sex on trabecular change overtime. Individuals with grossly observable mid-thoracic pathologies or trauma were excluded from analysis. The left 6^th^ rib was prioritized for consistency with traditional 2D histomorphometric age-at-death estimation methods. If the 6^th^ rib was not available, the 5^th^ or 7^th^ rib was used. If none of the left ribs were available, the right side was chosen following the same priority.

In preparation for CT analysis, the curve length of each rib was measured using a soft measuring tape along the cutaneous surface. Plastic spherical markers were placed at 50 and 75% of the total length from the vertebral end using hot glue adhesive. The plastic markers appear opaque in the CT images, allowing for region of interest (ROI) selection at precise locations (midshaft and anterior rib) in the post-processing stage of scanning. In groups of four, the ribs were placed in a custom-made foam fixture for scanning and both regions were imaged using a Northstar Inc. X5000 high-resolution computed tomography system with resolutions ranging from 35–49 microns, depending on the size of the ribs. Unlike other skeletal elements typically used in CT analysis, the unique curvature of each rib made it difficult to create a standard scanning position across all specimens. The use of the spherical markers and the custom designed foam fixture, which permitted for rib adjustment with each scan group, allowed for as much consistency as possible in scanning.

After the scans were completed, proprietary software for the Northstar X5000 was used for post-processing and extraction of the ROIs. Six-millimeter ROIs were selected sternally to each marker and processed using Dragonfly V4.1 (Figure 1). Dragonfly V4.1 was used to isolate cortical bone volumes of interest (VOIs) and three trabecular VOIs; one each along the cutaneous cortex, the center of the medullary cavity, and the pleural cortex (Figure1). Dragonfly V4.1 was also used to analyze cortical bone VOIs from the same 6mm sections sternal to the markers at the midshaft and anterior ROIs for average cortical area and total area. Segmentation of cortical and trabecular bone was accomplished using Buie segmentation (21) in the Bone Analysis tool.

**Fig. 1.**
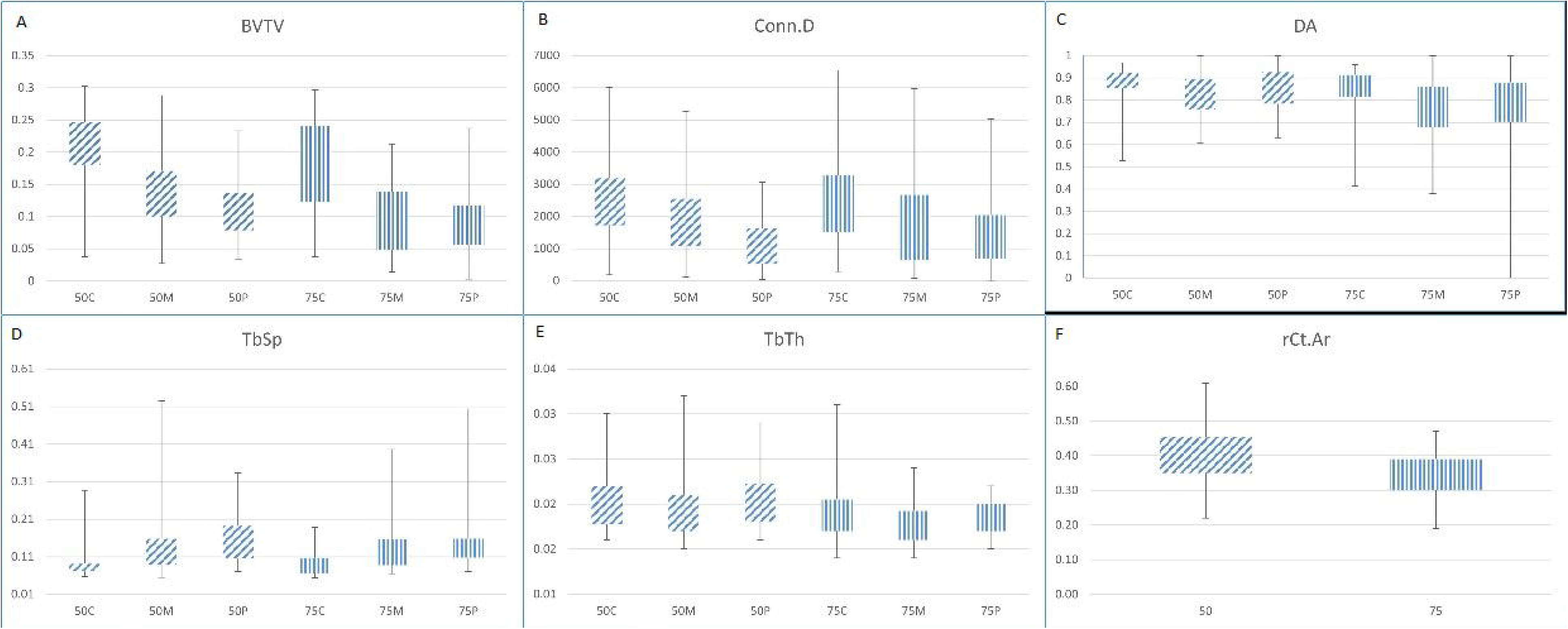
Illustration of region of interest (ROI) and volume of interest (VOI) selection. ROI was taken 6mm sternal to the marker. VOIs were selected by evenly dividing the medullary space into three volumes shown above: cutaneous (purple), medullary (green), and pleural (red).

Using the BoneJ plugin for ImageJ (22), each trabecular VOI was analyzed for bone volume fraction (BV/TV), trabecular thickness (TbTh), trabecular spacing (TbSp), connectivity density (Conn.D), and degree of anisotropy (DA), within and between the 50 and 75% ROIs (see Table 1). Independent sample *t*-tests, Kruskal-Wallis, or Mann-Whitney U tests were used to compare both trabecular and cortical variables within and between regions dependent on normality and groupings. The alpha values were set at 0.05 for all analyses.

**Table 1.**
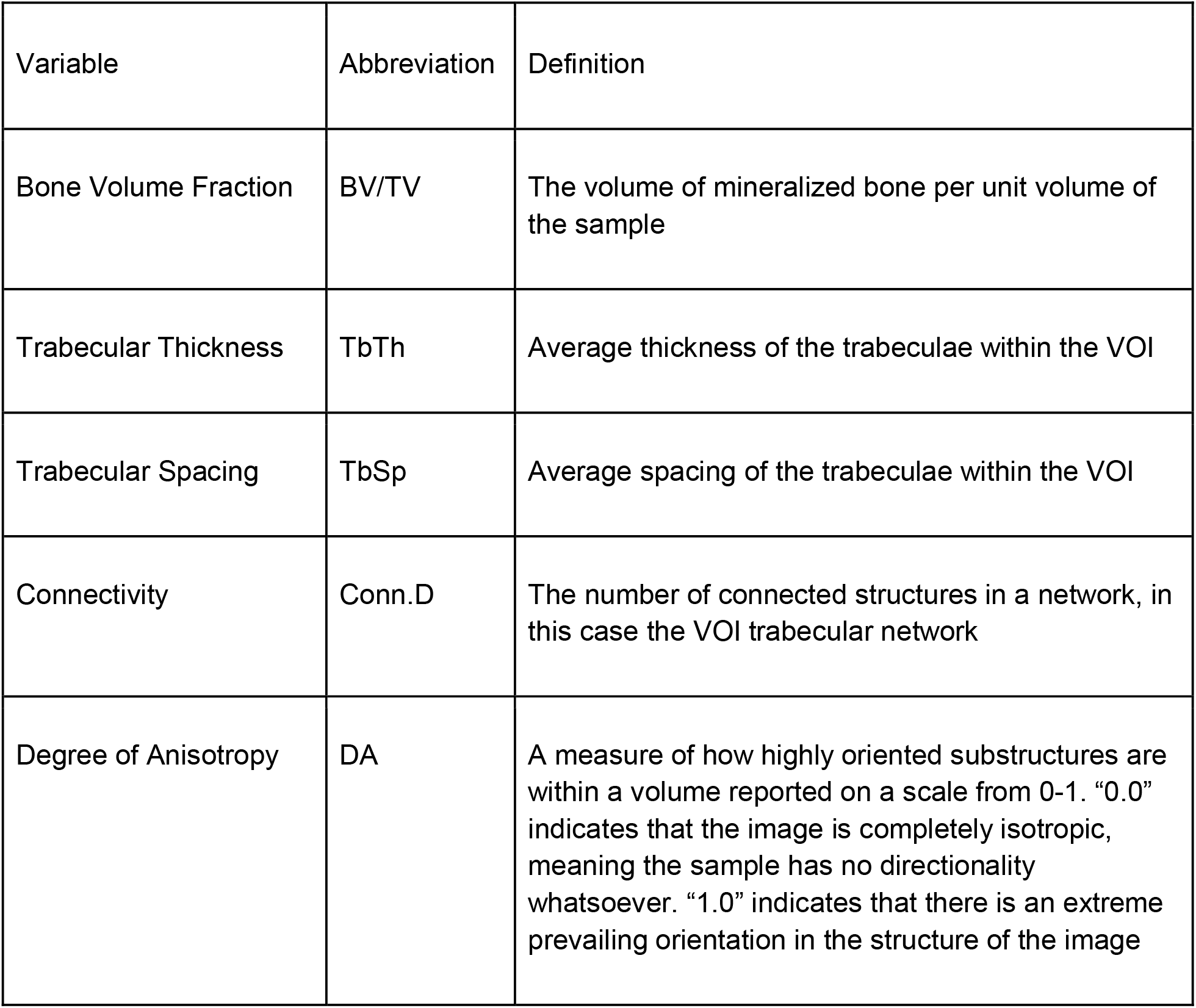
Trabecular variable definitions derived from BoneJ (22)

| Variable | Abbreviation | Definition |
| --- | --- | --- |
| Bone Volume Fraction | BV/TV | The volume of mineralized bone per unit volume of the sample |
| Trabecular Thickness | TbTh | Average thickness of the trabeculae within the VOI |
| Trabecular Spacing | TbSp | Average spacing of the trabeculae within the VOI |
| Connectivity | Conn.D | The number of connected structures in a network, in this case the VOI trabecular network |
| Degree of Anisotropy | DA | A measure of how highly oriented substructures are within a volume reported on a scale from 0-1. "0.0" indicates that the image is completely isotropic, meaning the sample has no directionality whatsoever. "1.0" indicates that there is an extreme prevailing orientation in the structure of the image |

## Results and Discussion

### Trabecular Analysis

Within both the 50 and 75% regions, cutaneous VOIs had significantly more BV/TV than the medullary and pleural VOIs. There was significantly greater Conn.D for the cutaneous VOIs than pleural VOIs in both regions, while there was significantly less TbSp in the cutaneous VOIs than either the medullary or pleural VOIs for both regions. These results show that in both ROIs the cutaneous portion of the rib contained significantly more bone that presented thicker struts with less space between struts (Figure 3). While detailed literature on the biomechanics of individual ribs throughout their length and *in situ* in the thorax is limited, these findings are consistent with preliminary work that indicates that the cutaneous and pleural surfaces of the ribs experience different strain modes during ventilation (23). Cutaneously, anterior and midshaft locations of the ribs undergo compression, while on the pleural surface these regions experience tension (23). This larger, thicker, and less spaced trabecular bone in the cutaneous VOIs may be an adaptation to these compressive loads.

**Fig. 2.**
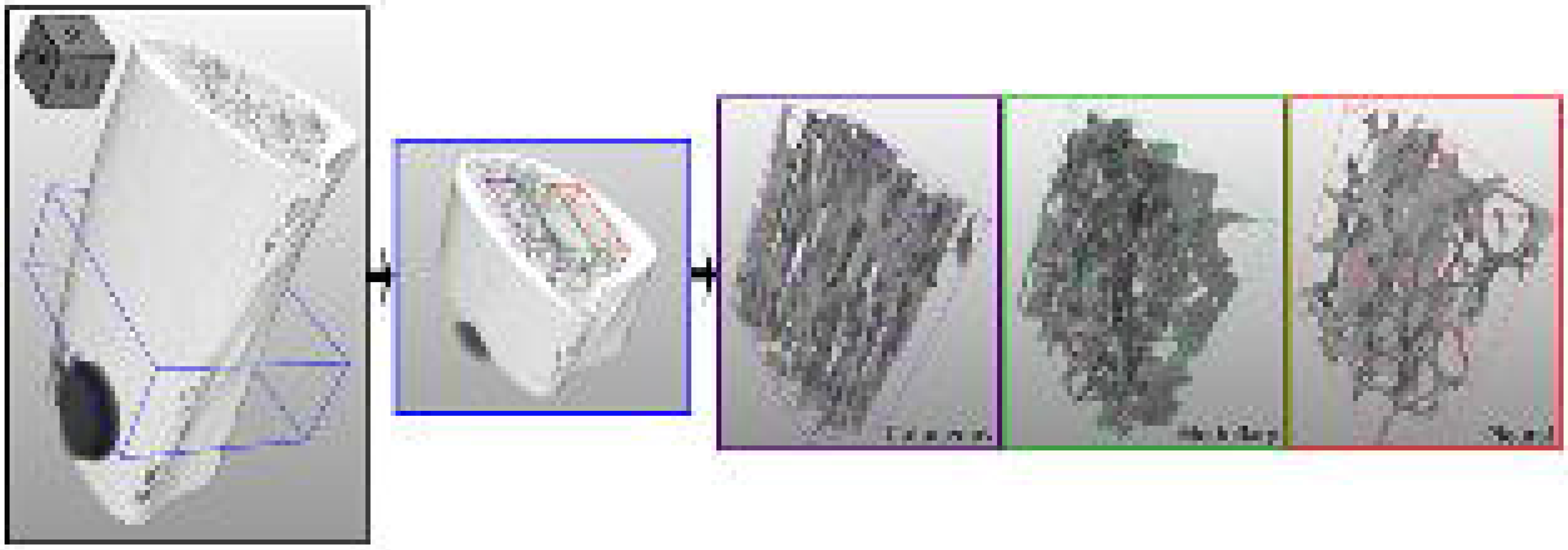
2D cross-sectional image of a rib at 50% length from different ages across the sample range, illustrating the 2D trabecular bone volume decrease with age. Images are to scale.

**Fig. 3.**
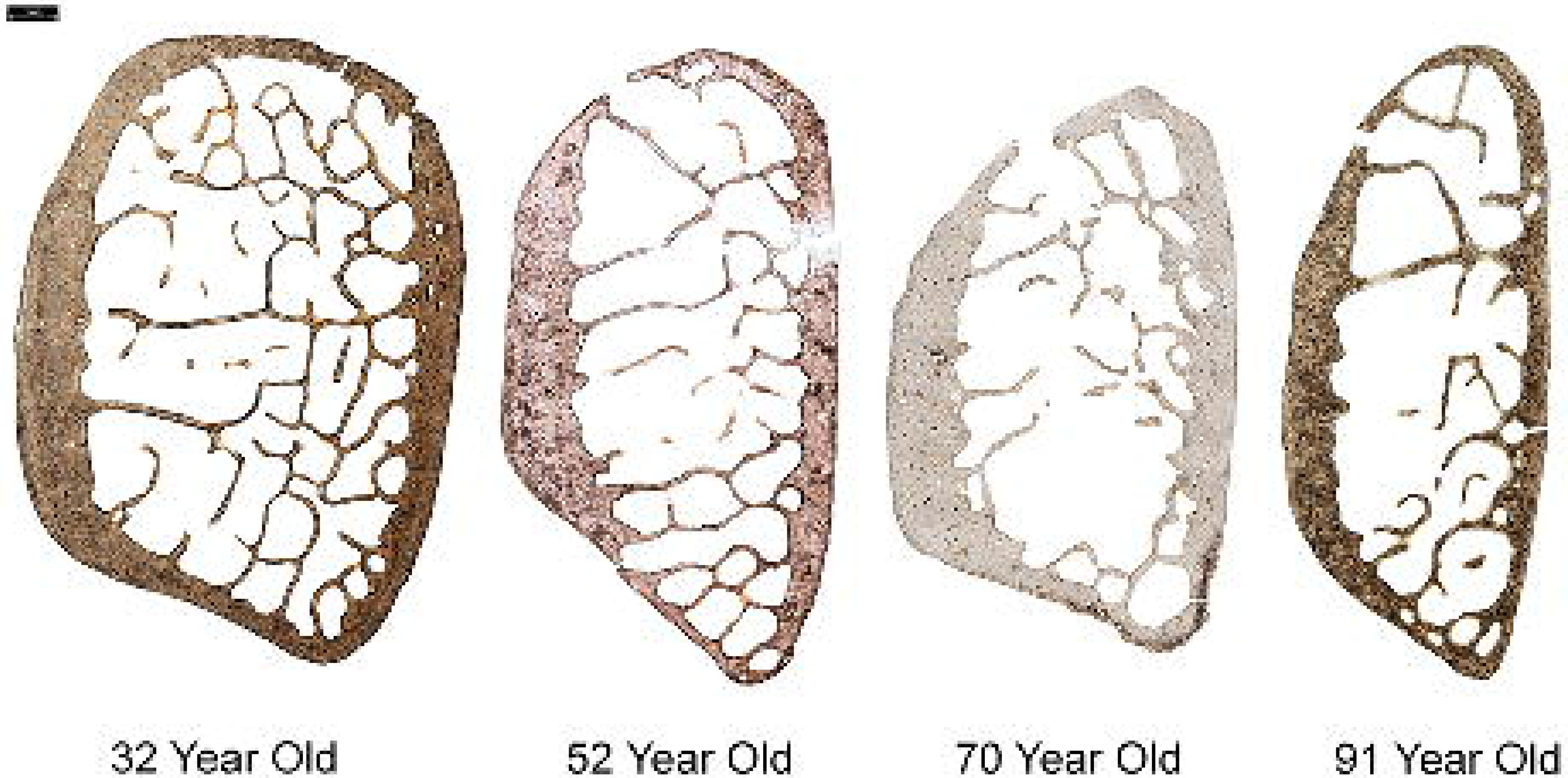
Boxplot distributions of trabecular variables within both 50 and 75% ROIs. Boxes A-E show the variables distributed across the cutaneous (C), medullary (M), and pleural (P) VOIs. A) Bone volume fraction (BV/TV); B) Connectivity density (Conn.D); C) Degree of anisotropy (DA); D) Average space between trabeculae (TbSp); E) Average trabecular thickness (TbTh); F) Relative cortical area (rCt.Ar) at the 50% and 75% ROIs.

Between the 50% and 75% ROIs, the 75% ROI showed more variation in BV/TV, DA, and TbTh overall, while the midshaft ROI had significantly more BV/TV, thicker trabeculae, and greater DA compared to the 75% region (Figure 3). The 50% ROI is thought to experience more consistent force than the 75% ROI by virtue of its position along the rib. In the midthoracic true ribs, the midshaft of the rib elevates during inspiration in what is described as a bucket handle movement, with rotation occurring at the posterior and anterior attachments of the thorax (24). The 75% ROI, attached to the sternum via costal cartilage and closer to that pivot point, experiences less consistent mechanical loading, leading to more variation in bone volume, orientation, and thickness. The individual VOIs showed no significant differences between regions in any variables.

The regression analysis used to examine the relationship of the trabecular variables (Table 2) with age show that BV/TV has weak but significant correlations with age at both the 50% (R^2^=0.041) and 75% (R^2^=0.068) regions. TbTh showed weak but significant correlations with age at the 75% ROI (R^2^=0.253), while TbSp showed weak but significant correlations with age at the 50% ROI (R^2^=0.040). There were no significant correlations between age and Conn.D or DA. The weak correlation with chronological age for all these variables suggest they will not significantly contribute to improved age estimation, although TbTh was the most highly correlated and may merit further investigation.

**Table 2.**
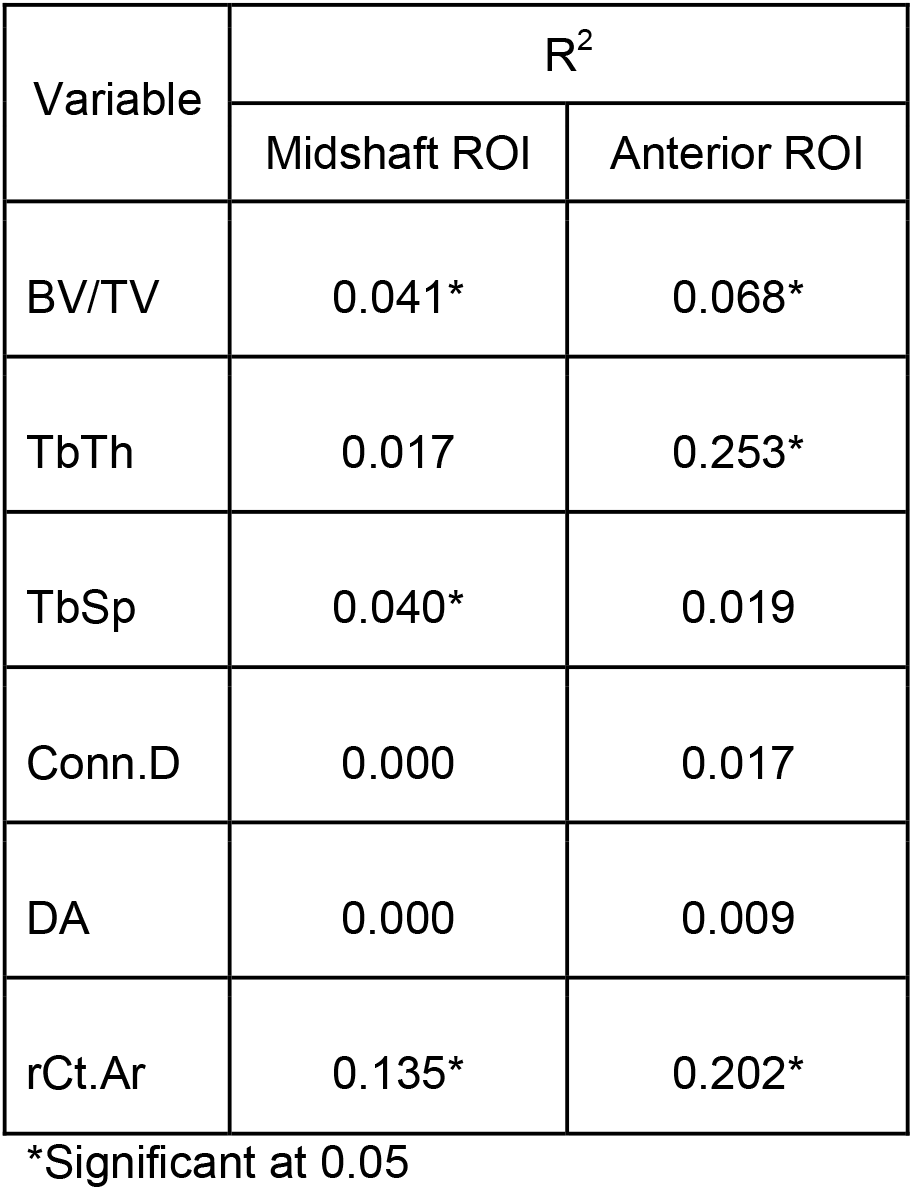
Correlation values of histological variables and age-at-death.

| Variable | R<sup>2</sup> |  |
| --- | --- | --- |
|  | Midshaft ROI | Anterior ROI |
| BV/TV | 0.041* | 0.068* |
| TbTh | 0.017 | 0.253* |
| TbSp | 0.040* | 0.019 |
| Conn.D | 0.000 | 0.017 |
| DA | 0.000 | 0.009 |
| rCt.Ar | 0.135* | 0.202* |

\*Significant at 0.05

### Cortical Analysis

Between the 50% and 75% ROIs, the cortical bone showed a significant difference in average relative cortical area (rCt.Ar) with the 50% region being significantly larger than the 75% ROI (Figure 4). When regressed against age, both the 50 and 75% regions showed significant but weak correlations with age (R^2^=0.135, R^2^=0.202) (Table2). The cortical results correlate with the trabecular results in that there is more bone in the 50% than the 75% ROIs, however, there is more variation in rCt.Ar within the 50% ROI than at 75% (Figure 3). While the R^2^ values for correlation between age and rCt.Ar area at both 50% and 75% are larger than the corresponding correlation values for BV/TV, both the BV/TV and rCt.Ar correlations are stronger with age at the 75% ROI than at the 50% ROI. Previous research by Dominguez et al. (25) has also shown significant differences between 50% and 75% ROIs in the rib, although this previous study examined total cortical area (Ct.Ar) rather than relative area. Since the Dominguez et al. (25) study found no significant difference in osteon population density (OPD) between the 50% and 75% ROIs, they suggest that when midshaft (50%) cannot be identified a more anterior, rather than a more posterior, section be taken for histological analysis. In light of the results of this study showing better correlation with age at the 75% region and the observation of no difference in OPD between midshaft and anterior ROIs from the previous study (25), using anterior rib sections for future histological age-at-death method development may produce adequate prediction models. However, more research should be done that includes pooled-sex samples to ensure the results agree with these findings.

Contrary to expectations, three-dimensional quantification of trabeculae did not elucidate a stronger relationship between trabecular bone loss and age than 2D analysis in the rib. Similar to previous 2D histological analysis of trabecular changes in the human rib (6, 17), there was a significant change in BV/TV with age, but the negative trend with age explains only 4-6% of the variation in bone volume. The most highly correlated variable with age was TbTh at the anterior position where age explained 25% of the variation in thickness.

Current methods for estimating skeletal age-at-death from rib histomorphometry are based on linear regression methods that make it difficult to surmount the problem of age mimicry and the problem of conditional dependence of histomorphometric variables (26). As the field moves towards more advanced statistical models for histological age estimation (9), it is possible that variables with low correlations with age could become useful contributors to age-at-death estimation methods when combined with other variables. The results of this study suggest that for male samples, 2D histological analysis of trabecular change is an appropriate method of trabecular analysis, and that 3D analysis does not greatly improve correlations between BV/TV and age-at-death. Future research should examine if the accelerated change with age seen in females (18) is better captured with 3D analysis.

## Acknowledgments

We would like to thank the donors whose gift made this research possible, the Society of Forensic Anthropologists for funding this research, and Devora Gleiber with data collection assistance. The Northstar X5000 was purchased through NSF MRI Award No.1338044 and the 2D images and analysis were done using equipment purchased through NSF MRI Award No. 1920218.

## Notes

### Competing Interest Statement

The authors have declared no competing interest.

## References

1. Stout SD, Paine RR. Brief communication: histological age estimation using rib and clavicle. Am J Phys Anthropol. 1992 Jan;87(1):111–5.10.1002/ajpa.1330870110

2. Cho H, Stout SD, Madsen RW, Streeter MA. Population-specific histological age-estimating method: a model for known African-American and European-American skeletal remains. J Forensic Sci. 2002 Jan;47(1):12–8

3. Kim YS, Kim DI, Park DK, Lee JH, Chung NE, Lee WT, et al. Assessment of histomorphological features of the sternal end of the fourth rib for age estimation in Koreans. J Forensic Sci. 2007 Nov;52(6):1237–42.10.1111/j.1556-4029.2007.00566.x

4. Pavon MV, Cucina A, Tiesler V. New formulas to estimate age at death in Maya populations using histomorphological changes in the fourth human rib*. J Forensic Sci. 2010 Mar 1;55(2):473–7.10.1111/j.1556-4029.2009.01265.x

5. Pfeiffer S, Heinrich J, Beresheim A, Alblas M. Cortical bone histomorphology of known-age skeletons from the Kirsten collection, Stellenbosch university, South Africa. Am J Phys Anthropol. 2016 May;160(1):137–47.10.1002/ajpa.22951

6. Andronowski JM, Crowder C. Bone Area Histomorphometry. J Forensic Sci. 2019 Mar;64(2):486–93.10.1111/1556-4029.13815

7. Cho H, Stout SD, Bishop TA. Cortical bone remodeling rates in a sample of African American and European American descent groups from the American Midwest: comparisons of age and sex in ribs. Am J Phys Anthropol. 2006 Jun;130(2):214–26.10.1002/ajpa.20312

8. Perz R, Toczyski J, Subit D. Variation in the human ribs geometrical properties and mechanical response based on X-ray computed tomography images resolution. J Mech Behav Biomed Mater. 2015 Jan;41:292–301.10.1016/j.jmbbm.2014.07.036

9. Kenyhercz MW, Crowder C, Dominguez VM, editors. Histological Age Estimation of the Femur Using Random Forest Regression. 72nd Annual Scientific Meeting of the American Academy of Forensic Sciences; 2020; Anaheim, CA.

10. Burr DB, Akkus O. Bone Morphology and Organization. In: Allen DBBR, editor. Basic and Applied Bone Biology. San Diego: Academic Press; 2014;3–25.

11. Frost HM. Bone “mass” and the “mechanostat”: a proposal. Anat Rec. 1987 Sep;219(1):1–9.10.1002/ar.1092190104

12. Kerley ER. The microscopic determination of age in human bone. Am J Phys Anthropol. 1965 Jun;23(2):149–63.10.1002/ajpa.1330230215

13. van Oers RF, Ruimerman R, Tanck E, Hilbers PA, Huiskes R. A unified theory for osteonal and hemi-osteonal remodeling. Bone. 2008 Feb;42(2):250–9.10.1016/j.bone.2007.10.009

14. Zebaze R, Seeman E. Cortical bone: a challenging geography. J Bone Miner Res. 2015 Jan;30(1):24–9.10.1002/jbmr.2419

15. Dominguez VM, Agnew AM. Examination of Factors Potentially Influencing Osteon Size in the Human Rib. Anatomical record. 2016 Mar;299(3):313–24.10.1002/ar.23305

16. Agnew AM, Stout SD. Brief communication: Reevaluating osteoporosis in human ribs: the role of intracortical porosity. Am J Phys Anthropol. 2012 Jul;148(3):462–6.10.1002/ajpa.22048

17. Beresheim AC, Pfeiffer SK, Alblas A. The Influence of Body Size and Bone Mass on Cortical Bone Histomorphometry in Human Ribs. Anatomical record. 2018 Oct;301(10):1788–96.10.1002/ar.23933

18. Beresheim AC, Pfeiffer SK, Grynpas MD, Alblas A. Sex-specific patterns in cortical and trabecular bone microstructure in the Kirsten Skeletal Collection, South Africa. American journal of human biology : the official journal of the Human Biology Council. 2018 May;30(3):e23108.10.1002/ajhb.23108

19. Treece GM, Gee AH, Mayhew PM, Poole KE. High resolution cortical bone thickness measurement from clinical CT data. Med Image Anal. 2010 Jun;14(3):276–90.10.1016/j.media.2010.01.003

20. Ryan TM, Walker A. Trabecular bone structure in the humeral and femoral heads of anthropoid primates. Anatomical record. 2010 Apr;293(4):719–29.10.1002/ar.21139

21. Buie HR, Campbell GM, Klinck RJ, MacNeil JA, Boyd SK. Automatic segmentation of cortical and trabecular compartments based on a dual threshold technique for in vivo micro-CT bone analysis. Bone. 2007 Oct;41(4):505–15.10.1016/j.bone.2007.07.007

22. Doube M, Klosowski MM, Arganda-Carreras I, Cordelieres FP, Dougherty RP, Jackson JS, et al. BoneJ: Free and extensible bone image analysis in ImageJ. Bone. 2010 Dec;47(6):1076–9.10.1016/j.bone.2010.08.023

23. Stemmer K. Characterization of Loading Environment on Human Ribs during Ventilation [Undergraduate Honors Thesis]: The Ohio State University, 2016.

24. Liebsch C, Wilke H-J. Chapter 3 – Basic Biomechanics of the Thoracic Spine and Rib Cage. In: Galbusera F, Wilke H-J, editors. Biomechanics of the Spine: Academic Press; 2018;35–50.

25. Dominguez VM, Harden AL, Wascher M, Agnew AM. Rib Variation at Multiple Locations and Implications for Histological Age Estimation*. Journal of Forensic Sciences. 2020 Nov;65(6):2108–11. https://doi.org/10.1111/1556-4029.14520

26. Konigsberg LW, Herrmann NP, Wescott DJ, Kimmerle EH. Estimation and Evidence in Forensic Anthropology: Age-at-Death. Journal of Forensic Sciences.2008 May;53(3):541–57. https://doi.org/10.1111/j.1556-4029.2008.00710.x

